# Whi5 hypo- and hyper-phosphorylation dynamics control cell cycle entry and progression

**DOI:** 10.1101/2023.11.02.565392

**Authors:** Jordan Xiao, Jonathan J. Turner, Mardo Kõivomägi, Jan M. Skotheim

## Abstract

Progression through the cell cycle depends on the phosphorylation of key substrates by cyclin-dependent kinases. In budding yeast, these substrates include the transcriptional inhibitor Whi5 that regulates the G1/S transition. In early G1 phase, Whi5 is hypo-phosphorylated and inhibits the SBF complex that promotes transcription of the cyclins *CLN1* and *CLN2*. In late-G1, Whi5 is rapidly hyper-phosphorylated by Cln1,2 in complex with the cyclin-dependent kinase Cdk1. This hyper-phosphorylation inactivates Whi5 and excludes it from the nucleus. Here, we set out to determine the molecular mechanisms responsible for Whi5’s multi-site phosphorylation and how they regulate the cell cycle. To do this, we first identified the 19 Whi5 sites that are appreciably phosphorylated and then determined which of these sites are responsible for G1 hypo-phosphorylation. Mutation of 7 sites removed G1 hypo-phosphorylation, increased cell size, and delayed the G1/S transition. Moreover, the rapidity of Whi5 hyper-phosphorylation in late G1 depends on ‘priming’ sites that dock the Cks1 subunit of Cln1,2-Cdk1 complexes. Hyper-phosphorylation is crucial for Whi5 nuclear export, normal cell size, full expression of SBF target genes, and timely progression through both the G1/S transition and S/G2/M phases. Thus, our work shows how Whi5 phosphorylation regulates the G1/S transition and how it is required for timely progression through S/G2/M phases and not only G1 as previously thought.

## INTRODUCTION

In many organisms, cells determine whether or not to divide by integrating diverse growth and differentiation signals at the G1/S transition ^1–3^. For example, cell growth drives progression through the G1/S transition in both budding yeast and human cells, while differentiation signals, such as those activated by mating pheromones in yeast, often inhibit the G1/S transition. In both budding yeast and human cells, the decision to divide is made irreversible by the activation of multiple positive feedback loops at the G1/S transition ^4–6^. After positive feedback activation, cells are relatively insensitive to changes in activities of the upstream growth and differentiation signals.

The integration of growth and differentiation signals at the G1/S transition is accomplished by the cell cycle control network ^6–9^. In yeast, multiple inputs regulate the activity of the central heterodimeric transcription factors SBF (Swi4/Swi6) and MBF (Mbp1/Swi6), which together control the expression of about 200 cell cycle-regulated genes responsible for diverse cellular functions including DNA replication and mitosis ^10–13^. SBF activity is inhibited by Whi5 and promoted by the G1 cyclin Cln3, most likely through direct protein-protein interactions ^8,14,15^. The irreversible decision to divide occurs when SBF activity increases enough to trigger a dramatic increase in the expression of the G1 cyclins *CLN1* and *CLN2* ^6^. Cln1,2-Cdk1 complexes then complete the phosphorylation of Whi5, which leads to its inactivation and exclusion from the nucleus. This further promotes SBF-dependent gene expression ^6,14^. SBF-dependent gene expression is also likely promoted by Cln1,2-Cdk1 phosphorylating SBF or SBF-bound proteins in addition to Whi5 ^16–18^. This series of positive feedbacks allows SBF to sustain its activity even when upstream signals such as Cln3 are removed ^4^.

While it was previously thought that Cln3-Cdk1 acted through Whi5 to activate SBF, it is now clear that Cln3 and Whi5 serve as separate inputs to SBF activity ^19^. This can be seen in the fact that Cln3 remains localized to SBF sites on the genome in *whi5*Δ cells ^19^. Moreover, cells expressing a hyperactive *CLN3* allele are smaller than *whi5*Δ cells and *cln3*Δ*whi5*Δ cells are larger than *whi5*Δ cells ^8,14,19^. Rather than target Whi5, as proposed in earlier models, Cln3-Cdk1 phosphorylates the C-terminal heptad repeats in RNAPII. In the revised model, Cln3-Cdk1 is recruited to SBF-regulated promoters as a direct transcriptional activator ^19^. Cln3 concentration likely reflects cellular growth rate as it is higher in rich nutrient conditions containing glucose than in poor nutrient conditions ^20^. In contrast, a similar amount of Whi5 protein is made in rich and poor nutrient conditions ^21^. Thus, Whi5 concentration directly reflects cell size prior to Start, even in different growth conditions, and is progressively diluted as cells grow in G1^21–25^. In this manner, Cln3 and Whi5 transmit distinct growth rate and cell size signals that are integrated at the G1/S transition.

While Whi5 concentration dynamics enable cells to sense their size in G1, Whi5 also exhibits a dynamic cell-cycle-dependent phosphorylation pattern that could transmit additional signals to the G1/S transition. In early G1, Whi5 exhibits diverse hypo-phosphorylated isoforms whose ratios remain approximately constant until the G1/S transition is initiated. Then, Whi5 is rapidly hyper-phosphorylated by Cln1,2-Cdk1 and exported from the nucleus ^16^ (**Fig. 1A**). While Whi5 hyper-phosphorylation and export from the nucleus leads to its inactivation, the function of the earlier hypo-phosphorylation was unknown. The early G1 Whi5 hypo-phosphorylation is not due to Cdk1 activity ^19^, suggesting that Whi5 integrates additional signals besides cell size in its role as an input to the G1/S decision. Alternatively, G1 hypo-phosphorylation could have little effect on the G1/S transition given that many of the isoforms are likely only mono-phosphorylated.

**Figure 1:**
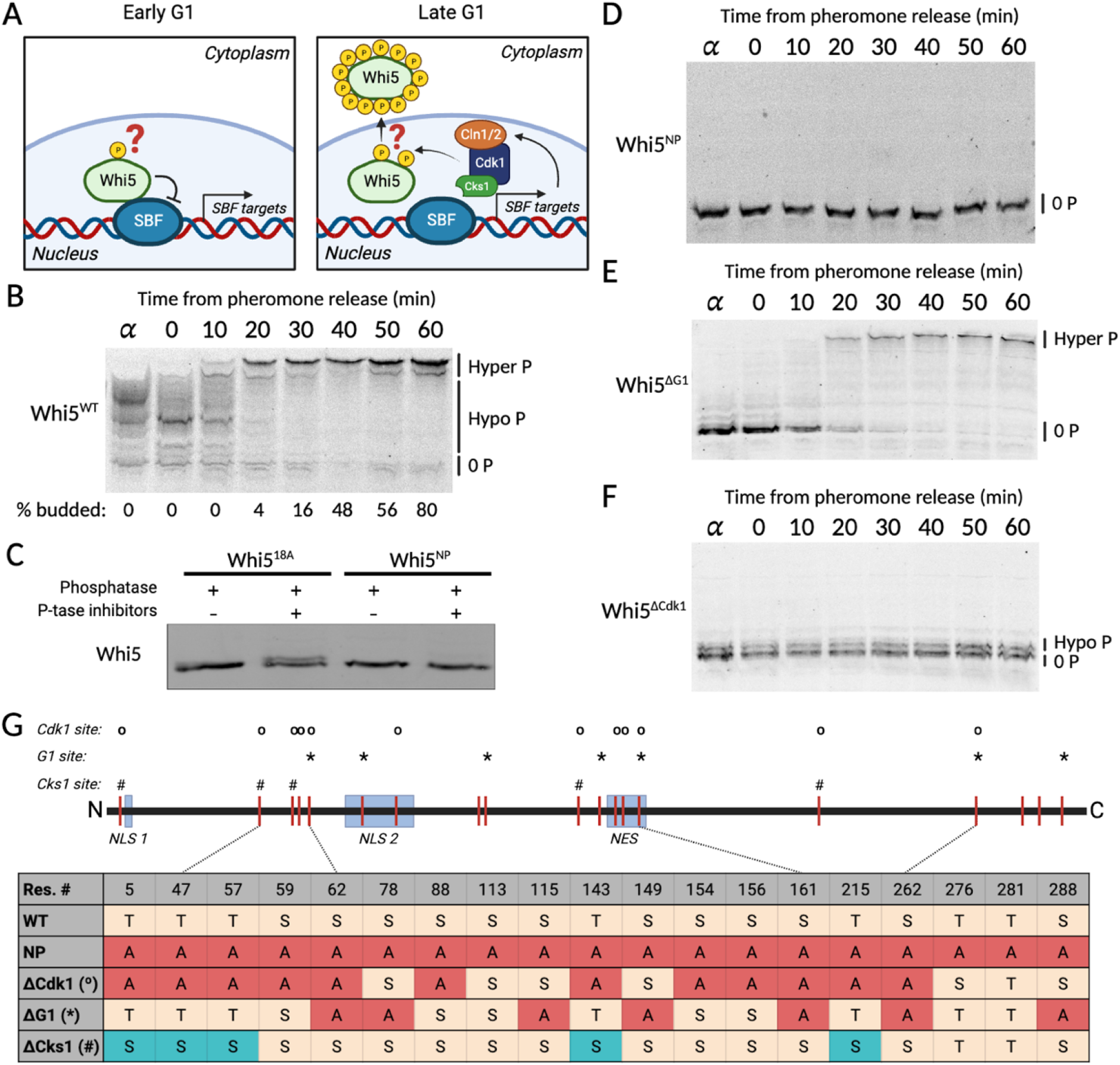
Identification of Whi5 phosphorylation sites. A) Schematic illustrating the dynamics of Whi5 phosphorylation through the G1/S transition. B) Phos-tag gel immunoblot of Whi5^WT^-3xFLAG. Cells were arrested in G1 using α-factor, then released into the cell cycle through drug washout. The percentage of cells budded at each timepoint is indicated at the bottom. C) Phos-tag gel immunoblot of Whi5^18A^-3xFLAG and its derivative Whi5^NP^-3xFLAG featuring an additional alanine substitution at serine 78. Cell lysates were prepared from asynchronous populations and treated with λ phosphatase in the presence or absence of phosphatase inhibitors as indicated. D-F) Phos-tag immunoblot time course for cells released from α-factor as with B, but for cells expressing Whi5^NP^-3xFLAG, Whi5^ΔG1^-3xFLAG, and Whi5^ΔCdk1^-3xFLAG. G) Summary of phosphorylation sites on Whi5 and substitutions made in the indicated variants.

Here, we sought to address the question of how Whi5 phosphorylation contributes to the G1/S transition. First, we identified the 7 sites phosphorylated in early G1: 3 Cdk sites and 4 non-Cdk sites. Mutation of these G1 sites to alanine increases cell size, but has no effect on Whi5 translocation and gene expression dynamics downstream of the G1/S transition. Moreover, we find that Cks1, a phospho-adaptor subunit of Cln1,2-Cdk1 complexes ^26,27^, is required for the rapid hyper-phosphorylation of Whi5 in late G1. Cks1 binds phosphothreonine residues on cyclin-Cdk substrates and enhances phosphorylation of other target sites on the same molecule ^26,27^. Mutation of Cdk1-targeted threonine residues to serines preserves phosphorylation on Whi5, but disrupts the semi-processive phosphorylation conferred by Cks1, and also leads to larger cells. This highlights the importance of Cks1-dependent hyper-phosphorylation for timely cell cycle progression. Lastly, we confirmed the importance of Cdk1-dependent phosphorylation for Whi5 nuclear export ^14,28,29^, as Whi5 variants lacking the 12 Cdk1 phosphorylation sites or all 19 phosphorylation sites remained nuclear throughout the cell cycle. Disrupted nuclear export corresponded to a decrease in expression of *CLN2* and prolonged S/G2/M phases, demonstrating the importance of Whi5 phosphorylation for relieving SBF inhibition. Taken together, our results demonstrate that both Cdk1-independent hypo-phosphorylation and Cdk1-dependent hyper-phosphorylation of Whi5 promotes cell cycle progression by advancing the G1/S transition and by maintaining timely progression through S/G2/M, respectively. This demonstrates how a single protein such as Whi5 can serve as a hub integrating multiple signals to control cell cycle progression.

## RESULTS

### Identification and characterization of Whi5 phosphorylation sites

To address the question of how Whi5 phosphorylation dynamics contributes to the G1/S transition, we first sought to identify where and when residues on Whi5 were phosphorylated. To do this, we used Phos-tag immunoblotting to separate diverse phospho-isoforms of Whi5 ^19,30^. Cells were sampled from populations synchronously progressing through the cell cycle following release from a G1 arrest imposed using mating pheromone. Wild-type Whi5 is initially hypo-phosphorylated in mating arrest. Upon removal of mating pheromone, this hypo-phosphorylation decreases slightly before Whi5 is rapidly hyper-phosphorylated at the G1/S transition, consistent with previous results ^19^ (**Fig. 1B**).

After re-establishing the phosphorylation dynamics of wild-type Whi5, we next sought to identify all the sites that were phosphorylated. To do this, we first examined the phosphorylation of a Whi5 variant Whi5^18A^, in which all 18 previously identified phosphorylation sites were substituted with non-phosphorylatable alanines ^16^. However, phosphatase treatment of Whi5^18A^ isolated from asynchronously growing cells removed the upper band on the Phos-tag gel (**Fig. 1C**), indicating the presence of at least one additional phosphorylated site. To identify this additional phosphorylated site, we systematically mutated other sites on Whi5 identified in proteomics data ^17^. We found that the additional alanine substitution of serine 78 produces a fully un-phosphorylatable Whi5 (Whi5^NP^) as indicated by the lack of a Phos-tag band shift following phosphatase treatment (**Fig. 1C**). Whi5^NP^ remained unphosphorylated throughout the cell cycle (**Fig. 1D**).

Having identified the Whi5 phosphorylation sites, we next sought to identify the subset of these sites that are phosphorylated throughout G1. To do this, we added back sites one at a time to Whi5^NP^ and looked for Whi5 variants that were phosphorylated in early G1. This identified 7 G1 phosphorylation sites. Mutation of all 7 G1 sites resulted in a Whi5 variant (Whi5^ΔG1^) that exhibited no phosphorylation in G1 (**Fig. 1E**), while still experiencing post-Start hyper-phosphorylation. This confirms that these 7 sites are responsible for the hypo-phosphorylation of Whi5 in G1. Importantly, it is the ability of Phos-tag gels to resolve different mono-phosphorylated isoforms of Whi5 that allowed us to identify which sites are phosphorylated in G1 and which sites are phosphorylated through the rest of the cell cycle. The difference from previous work identifying only 18 Whi5 sites ^16^ is due to the improved resolution of our Phos-tag methods.

In addition to identifying phosphorylated sites and their dynamics through the cell cycle, we also sought to disrupt distinct groups of phosphorylation sites. In particular, we examined the effects of disruption of the 12 minimal Cdk1 consensus sites (S/TP) identified in previous studies ^16^ (Whi5^ΔCdk1^). Whi5^ΔCdk1^ experienced no change in phosphorylation as cells progressed through the cell cycle (**Fig. 1F**), which is consistent with previous work indicating that Cln1/2-Cdk1 is primarily responsible for the S/G2/M hyper-phosphorylated state ^6,8,31–33^. The Whi5 phosphorylation sites identified in our Phos-tag analyses and previous studies, as well as disruptions made in various mutants used in this study, are summarized in **Figure 1G**. In summary, this analysis revealed which sites are phosphorylated in G1 and which are subsequently phosphorylated at the G1/S transition by Cdk1.

### Cks1 is necessary for rapid and complete hyper-phosphorylation of Whi5

That Cdk1 consensus phosphorylation sites on Whi5 are the only ones rapidly phosphorylated at the G1/S transition suggests that Cdk1 is the kinase exclusively responsible for the hyper-phosphorylation of Whi5. This is supported by previous experiments examining cells expressing an ATP-analog-sensitive version of Cdk1. When a population of cells past the G1/S transition with hyper-phosphorylated Whi5 were treated with an ATP analog to inhibit Cdk1, Whi5 was rapidly dephosphorylated ^19^.

As Cdk1 is responsible for Whi5 hyper-phosphorylation, we next sought to examine molecular mechanisms underlying these phosphorylation dynamics. Previous work showed that Whi5 is a specific target of Cln2-Cdk1 complexes ^31^ and that part of this specificity is due to an LP linear docking motif ^32^. However, comparing Whi5 to Sic1, another cell cycle inhibitor hyper-phosphorylated at the G1/S transition, suggests that additional mechanisms besides LP docking contribute to Whi5 hyper-phosphorylation dynamics because that is the case for Sic1 ^31^. One additional determinant of Sic1 hyper-phosphorylation at the G1/S transition is Cks1, an additional, small subunit present in many cyclin-Cdk complexes that docks to phosphorylated threonine residues ^26,27^ (**Fig. 2A**). Thus, the presence of TP sites on a substrate, as is the case for Whi5, is predicted to enhance multi-site phosphorylation by providing additional docking platforms for cyclin-Cdk1-Cks1 complexes. Consistent with the prediction that Whi5 hyper-phosphorylation depends on Cks1, hypo-phosphorylated Whi5 is converted to hyper-phosphorylated Whi5 in late G1 without the preceding appearance of many intermediate phospho-isoforms (**Fig. 1B**).

**Figure 2:**
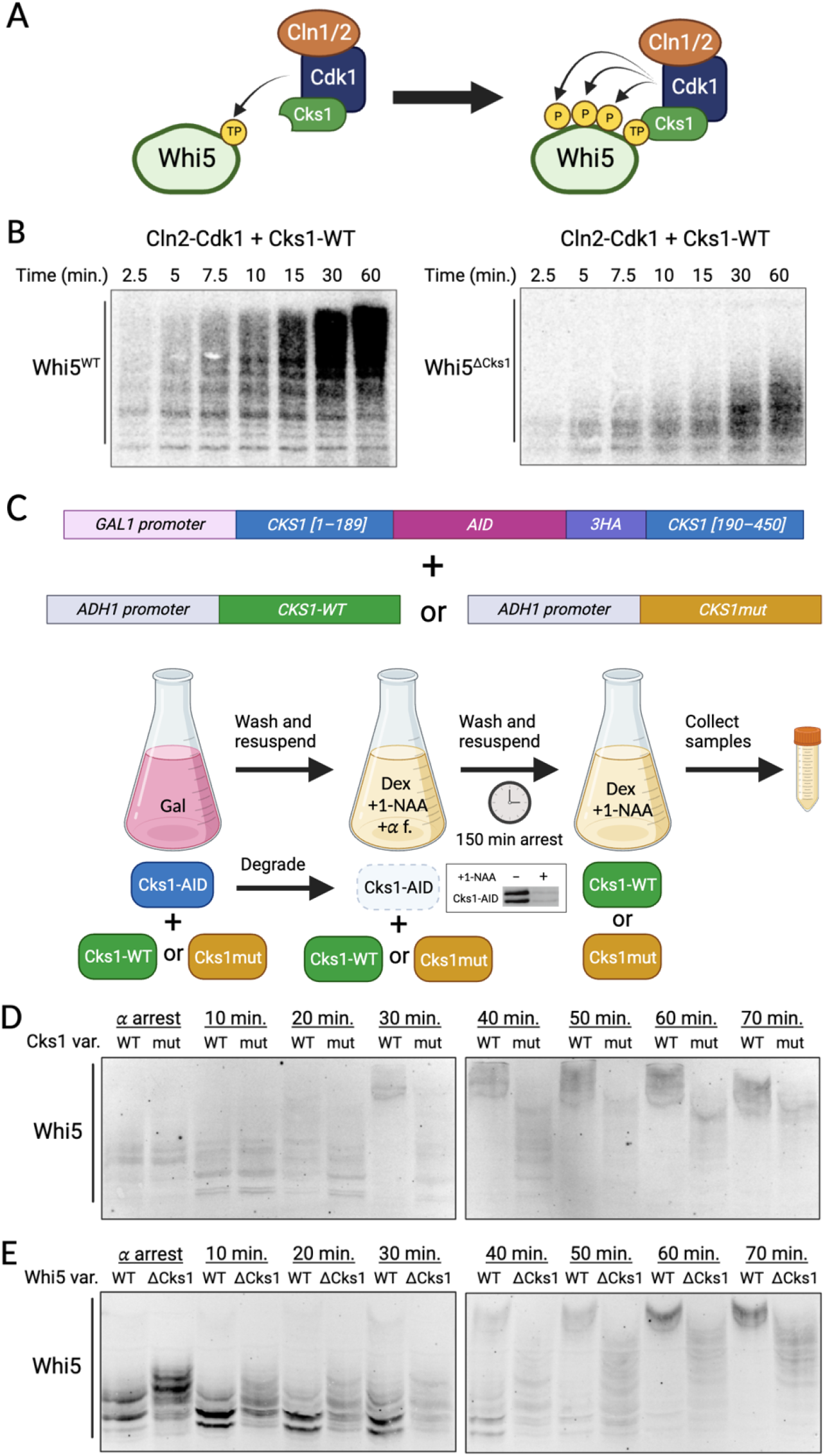
Cks1 contributes to the rapid hyper-phosphorylation of Whi5 during the G1/S transition. A) Schematic showing phospho-threonine docking of Cks1 driving Whi5 hyperphosphorylation. B) Autoradiograph showing the results from radiolabeled kinase assays using Cln2-Cdk1-Cks1 and either wild type Whi5 or a variant lacking TP residues that cannot be docked by Cks1 (Whi5^ΔCks1^). C) Schematic representation of the reagents and the experiment in D. *CKS1* was placed under control of the galactose-regulated promoter *GAL1*, and the protein was made conditionally unstable by inserting an auxin-inducible degron (AID) into an unstructured domain in the protein. Switching to glucose media and addition of +1-NAA shuts off synthesis of Cks1-AID and activates degradation of existing Cks1-AID. Inset shows immunoblot of Cks1-AID-3HA with and without +1-NAA. Cells were arrested in G1 using α-factor and then released into the cell cycle. D) Phos-tag immunoblot of Whi5^WT^-3xFLAG in mating factor arrest-and-release time course as described in C. E) Phos-tag immunoblot of Whi5^WT^-3xFLAG or Whi5^ΔCks1^-3xFLAG in otherwise wild-type cells. Cells were arrested in G1 using α-factor and then released into the cell cycle.

To test if Cks1-dependent docking contributed to Whi5 hyper-phosphorylation, we first examined the phosphorylation of Whi5 by Cln2-Cdk1-Cks1 complexes *in vitro*. To do this, we purified Cln2-Cdk1 complexes from yeast and then added Cks1 and Whi5 purified from bacteria to a reaction containing γ-32P ATP. The resulting Whi5 phospho-isoforms were analyzed using Phos-tag SDS-PAGE gels and autoradiography. The pattern of Whi5^WT^ phosphorylation by Cln2-Cdk1-Cks1 was consistent with semi-processive phosphorylation, in that hyper-phosphorylated species appeared and increased simultaneously with intermediately-phosphorylated species (**Fig. 2B**). Since Cks1 is known to bind exclusively to phosphothreonine residues, we also purified and phosphorylated a Whi5 variant in which all the TP sites were substituted with SP sites (Whi5^ΔCks1^, **Fig. 1G**). Phosphorylation of Whi5^ΔCks1^ by Cln2-Cdk1-Cks1 was substantially slower than the phosphorylation of Whi5^WT^ (**Fig. 2B**). These *in vitro* results support the hypothesis that rapid, semi-processive, hyper-phosphorylation of Whi5 depends on Cks1 docking to phosphothreonine residues.

Having demonstrated the importance of Cks1 for hyper-phosphorylation of Whi5 *in vitro*, we next sought to test the importance of Cks1 for Whi5 phosphorylation *in vivo*. To selectively deplete Cks1 from cells during pheromone-synchronized time courses, we constructed a version of *CKS1* fused to an auxin-inducible degron (Cks1-AID) and expressed it under the *GAL1* promoter. This construct replaced the endogenous *CKS1* gene (**Fig. 2C**). We were thus able to grow cells in galactose-containing media and then transfer them to glucose- and auxin-containing media to deplete Cks1-AID during and after mating arrest. To isolate the effects of Cks1 phosphothreonine binding, we did not simply deplete Cks1-AID but replaced it with either wild-type Cks1 or a variant (Cks1mut) with substitutions of three key residues in its phospho-threonine binding pocket (R33E/S82E/R102A) (**Fig. 2C**). Compared to the wild-type Cks1-expressing control cells, cells expressing Cks1mut phosphorylated Whi5 more slowly (**Fig. 2D**). This supports the view that Cks1-dependent priming contributes to the rapid accumulation of hyper-phosphorylated Whi5 *in vivo*. To confirm that the slower Whi5 phosphorylation observed in cells expressing Cks1mut was a direct effect of disrupting the Cks1-Whi5 interaction, rather than a consequence of disrupting a different Cks1-substrate interaction, we observed the phosphorylation dynamics of Whi5^ΔCks1^ in otherwise-wild-type cells. Whi5^ΔCks1^ was gradually phosphorylated following release from mating arrest (**Fig. 2E**), similar to Whi5^WT^ in cells expressing Cks1mut. This was true despite an increase in basal phosphorylation during G1, likely a result of a reduced dephosphorylation rate for phospho-serines compared to phospho-threonines ^34^. Taken together, these results indicate that rapid Whi5 phosphorylation relies on Cks1 binding to phosphothreonines on Whi5.

### Cdk1 phosphorylation of Whi5 drives nuclear export

Having defined sets of phosphorylation sites whose mutations target distinct aspects of Whi5 phosphoregulation, we next sought to determine how each set of sites affects the localization of Whi5, a well-established component of its cell cycle regulation ^14,16,28,29,35^. In late G1, hyperphosphorylation of Whi5 by Cln1,2-Cdk1 induces Whi5 dissociation from the SBF transcription factor and nuclear export.

To test how our mutations affect Whi5 localization dynamics, we fused the fluorophore mCitrine to the C-terminus of different variants of *WHI5*. It was previously shown that fusing mCitrine to the C-terminus does not affect Whi5 activity, since cell cycle progression and cell size are similar in Whi5-mCitrine and wild type cells ^22^. Whi5^WT^ is exported early in the G1/S transition, and remains in the cytoplasm until shortly before mitosis, when it is reimported and partitioned between the nuclei of the mother and daughter cells ^23^. This is apparent by both visual inspection and quantification using the coefficient of variation of fluorescence intensity over the cell area as a proxy for nuclear localization (**Fig. 3A,B**). In contrast, Whi5 variants lacking Cdk1-dependent phosphorylation sites (*i.e.* Whi5^NP^, Whi5^ΔCdk1^) show nuclear localization throughout the cell cycle. This is consistent with previous data showing that Whi5 nuclear export was inhibited by alanine substitution at 3 sites (Ser154, Ser156, and/or Ser161) in one nuclear export sequence ^28,29^ (**Fig. 1G**). Meanwhile, variants where the NES is left intact (*e.g.*, Whi5^ΔCks1^) show the same export behavior as Whi5^WT^. Interestingly, Whi5^ΔG1^ shows a ∼12-minute delay in nuclear export relative to Whi5^WT^, possibly due to one of the three Cdk1-target phosphorylation sites being mutated to alanine (Ser161). This suggests that SBF-dependent transcription driving bud emergence is activated prior to exporting Whi5.

**Figure 3:**
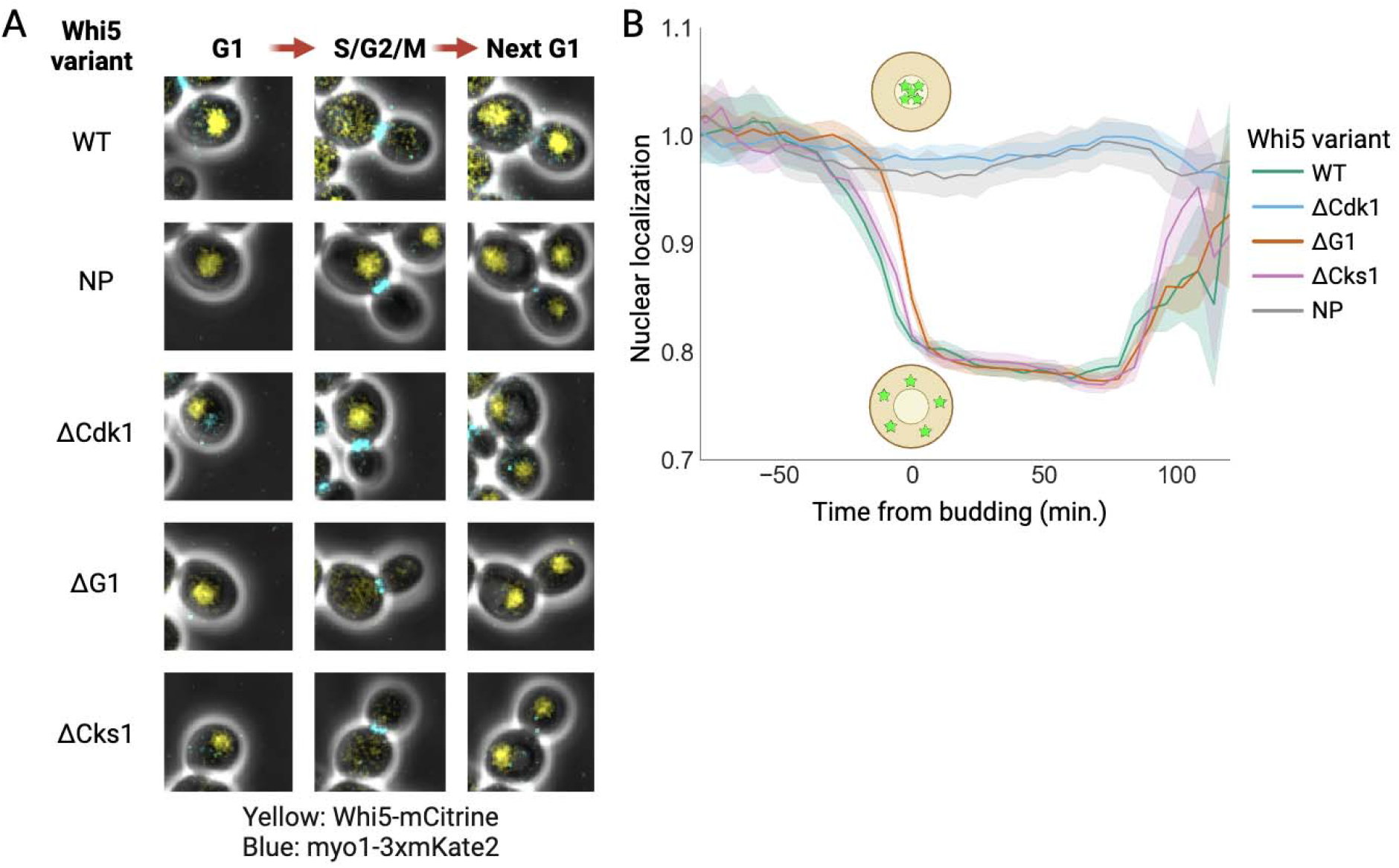
Whi5 localization dynamics. A) Composite phase and fluorescence images showing localization through the cell cycle for the indicated Whi5 variant fused to mCitrine. The transition from G1 to S/G2/M was determined by visual detection of a bud, while the transition from S/G2/M to the next G1 was determined by the loss of an actomyosin-mKate2 signal from the bud neck. Cells are grown in synthetic complete media with glycerol/ethanol as carbon sources. B) Averaged nuclear Whi5 signal from many single cell traces for indicated Whi5 variants (N_WT_ = 100, N_ΔCdk1_ = 200, N_ΔG1_ = 217, N_NP_ = 83, N_ΔCks1_ = 104). Traces are aligned at budding. The nuclear localization metric is the coefficient of variation of the Whi5-mCitrine fluorescence intensity over the cell area rescaled to its value in early G1. Data lines indicate the mean values and the shaded area denotes 95% c.i.

### Whi5 phosphorylation on G1 and Cdk1 sites regulates cell size

Next, we sought to determine how the phosphorylation of Whi5 affects its synthesis pattern, which was previously shown to regulate cell size. Whi5^WT^ was shown to be synthesized mostly during S/G2/M phases and then diluted through G1. The dilution of Whi5 in G1 enables cells to sense their size and is thus an important mechanism through which cell growth triggers the G1/S transition ^21–23,25^. Importantly, the amount of Whi5 synthesized in S/G2/M is largely independent of cell size, and Whi5 is partitioned in nearly equal amounts to the mother and daughter cells ^23^. This causes the concentration of Whi5 in daughter cells to be roughly inversely proportional to their size.

To test the effect of phosphorylation sites on Whi5 synthesis, we tracked asynchronously dividing cells expressing Whi5 variants fused to mCitrine. We restricted our analysis to the first cell cycle of first-generation daughter cells, which exhibit cell size control through an extensive G1 phase ^36–38^. These cells are therefore most affected by Whi5’s role as a size-sensing cell cycle regulator. Consistent with previous results ^21–23,25^, we observed that Whi5^WT^ is minimally synthesized in G1. This results in a nearly constant total protein amount until the G1/S transition, following which Whi5 is synthesized such that the total protein amount at mitosis is approximately double that at the start of the cell cycle (**Fig. 4A,B**). All Whi5 phosphorylation site mutants under the *WHI5* promoter show the same pattern of synthesis over the cell cycle, but strains lacking Cdk1 sites on Whi5 (*e.g.*, Whi5^NP^, Whi5^ΔCdk1^) both show reduced amounts of Whi5 protein at all points in the cell cycle (**Fig. 4A**). Interestingly, Whi5^ΔCdk1^ shows a nearly 50% reduction in total Whi5 protein relative to Whi5^WT^ even though their population doubling times were similar (∼150 minutes each). To test if the reduced Whi5^ΔCdk1^ amounts were due to a change in protein stability or synthesis, we created a construct using a drug-inducible promoter ^39^ to express and subsequently shut off Whi5-mCitrine expression to monitor its decay. Whi5^WT^ and Whi5^ΔCdk1^ both showed half lives >300 minutes (**Fig. S1**), which is much longer than the ∼180 minutes it takes for a cell to double its biomass when growing on ethanol as a carbon source. Thus, mutation of the Cdk1 sites on Whi5 reduces its synthesis, which can be compensated for by introducing a second copy of *WHI5pr-WHI5*^Δ*Cdk1*^.

**Figure 4:**
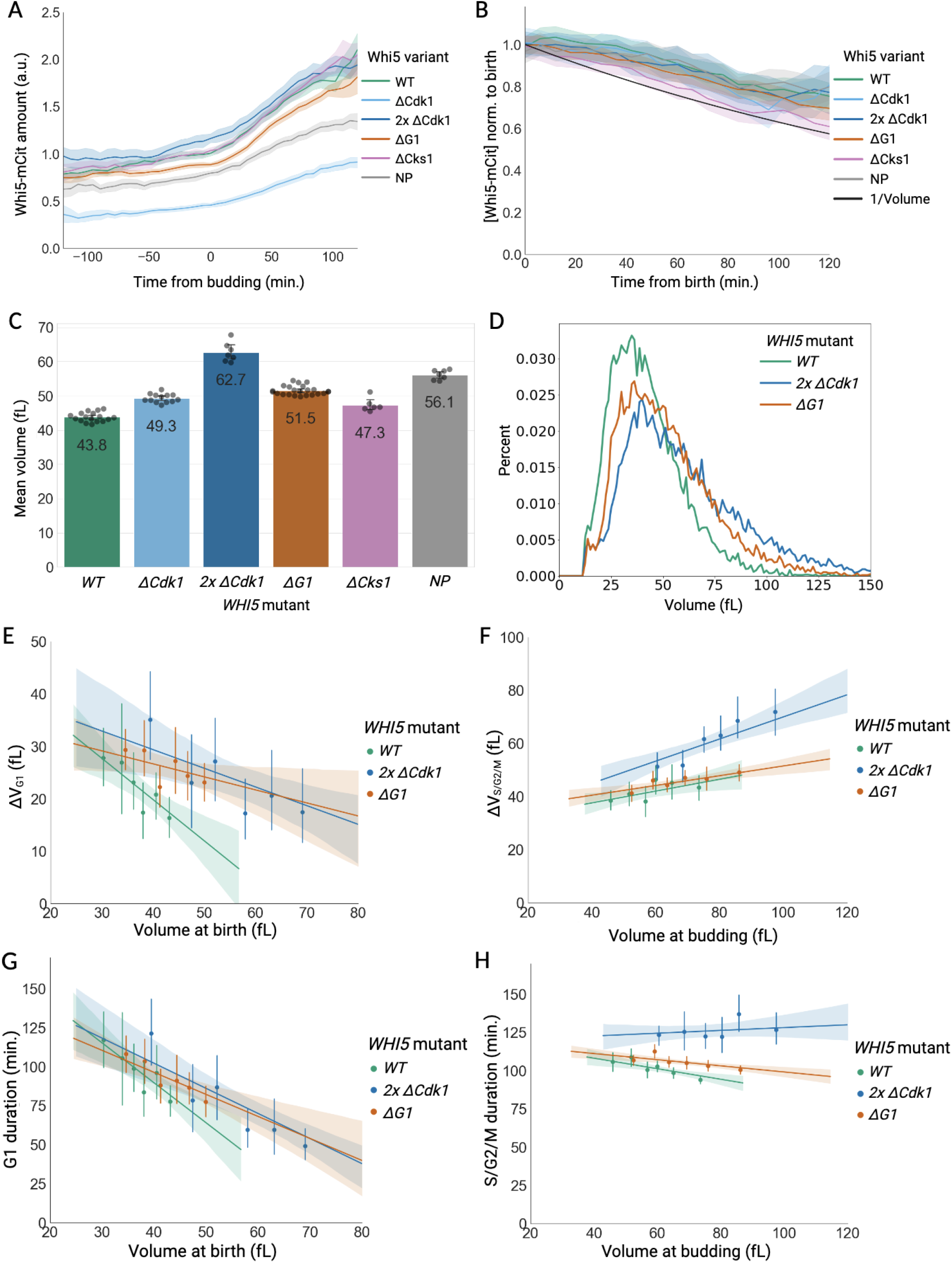
Whi5 synthesis dynamics and its effect on cell size control. A) Total Whi5-mCitrine signal averaged over many daughter cell cycles during growth in synthetic complete media with glycerol/ethanol as the carbon sources (population doubling time ∼150 minutes). Cells use a *WHI5* promoter to express the indicated Whi5 variant fused to mCitrine. Traces are aligned at bud appearance, and fluorescence was scaled so that WT cells expressed 1 a.u. at budding. Birth timing is determined by loss of an actomyosin-mKate2 signal from the bud neck. Data lines indicate the mean and the shaded area denotes the 95% c.i. N_WT_ = 100, N_ΔCdk1_ = 200, N_2x ΔCdk1_ = 98, N_ΔG1_ = 217, N_ΔCks1_ = 104, N_NP_ = 83. Data for all strains except *2xWHI5*^Δ*Cdk1*^ is from the same experiments as Figure 3. B) Average Whi5-mCitrine concentration aligned to the time of birth and normalized to the concentration at birth for each averaged trace shows dilution through G1. Perfect dilution, *i.e.*, zero synthesis and exponential volume increase over time (one doubling over 150 minutes), is shown in black. Same cells as in panel A. C) Mean volume of asynchronously growing cultures measured by Coulter counter. Bar height denotes mean, and error bars denote 95% c.i. D) Representative Coulter counter cell size distributions for the indicated *WHI5* genotype. E) Volume growth from birth to budding versus volume at birth for the same cells as panels A, B. Cells are separated into bins with equal numbers. Dots indicate the mean values per bin and error bars denote the 95% c.i. Solid line indicates linear regression to unbinned data and shaded areas denote 95% c.i. F) Volume growth between budding and mitosis versus volume at budding for the same cells as panels A, B. G) Duration of G1 phase versus volume at birth for the same cells as panels A, B. H) Duration of S/G2/M phases versus volume at budding for the same cells as panels A, B. The same plots as E-H with all genotypes are shown in Fig. S2C-F.

Having generated a series of strains expressing Whi5 variants at comparable amounts, we then assessed the ability of these proteins to regulate cell size (**Fig. 4C,D**). That *2xWHI5*^Δ*Cdk1*^ cells are much larger than *WHI5^WT^*cells even though both have a similar amount of Whi5 protein shows the importance of Cdk1 phosphorylation for G1/S regulation. We further observed that cells expressing Whi5^ΔCks1^ are slightly larger than Whi5^WT^, showing that the size-regulating function of Cdk1-dependent phosphorylation is partially dependent on the rapid hyper-phosphorylation afforded by Cks1-dependent docking. Lastly, cells expressing Whi5^ΔG1^ exhibit a ∼15% size increase compared to wild-type cells, which shows the importance of Whi5 phosphoregulation in G1. These data show that all phosphorylation site mutations that we tested make Whi5 a more potent cell cycle inhibitor. It is of particular interest that Whi5^ΔG1^ shows a striking size effect even though its nuclear export is intact, albeit delayed.

After establishing the effects of Whi5 phosphorylation site mutants on cell size, we next sought to test which parts of the cell cycle were affected in each case. To do this, we first used time lapse imaging to measure G1 size regulation in daughter cells. Wild-type cells exhibited a negative correlation between cell size at birth and the amount of volume growth and time spent in G1, similar to previously published results ^40^. *2xWHI5*^Δ*Cdk1*^ and *WHI5*^Δ*G1*^ daughter cells exhibited a similar negative correlation between cell size at birth and growth in G1, but both mutant cells grew more and spent longer in G1 than WT cells when cells were initially the same size at birth (**Fig. 4E,G**). These mutations did not affect the growth rate per unit mass during G1 (**Fig. S2A**). Interestingly, we found that mutation of Cdk1 phosphorylation sites also affected the duration of S/G2/M phases (**Fig. 4F,H**), which also affects average cell size ^41^. The effect on S/G2/M duration is only present in *2xWHI5*^Δ*Cdk*1^ rather than *WHI5*^Δ*G1*^ cells, which could still be phosphorylated by Cdk1 on some sites and exported from the nucleus. Interestingly, cells experience roughly the same extension of S/G2/M with one or two copies of *WHI5*^Δ*Cdk1*^, suggesting the effect is saturated (**Fig. S2F)**. We refer the reader to the supporting material for a similar single cell growth analysis of all genotypes (**Fig. S2-3**). Taken together our data indicates that Whi5 phosphorylation is important for both progression through G1 and S/G2/M phases.

### *CLN2pr* expression dynamics and transcription of SBF target genes

The cell cycle defects in the S/G2/M phases of cells expressing Whi5^ΔCdk1^ point to the importance of Cdk1 phosphorylation of Whi5 in late G1 for subsequent cell cycle events. This defect could be due to insufficient expression of the approximately 100 target genes regulated by Whi5’s main target, SBF ^11,15,19,42^. Whi5 proteins lacking Cdk1 sites also remain nuclear throughout the cell cycle, which would be expected to enhance its inhibitory activity. We therefore sought to investigate the effects of different Whi5 phosphorylation site mutations on SBF-dependent gene expression.

To test the effect of Whi5 phosphorylation sites on SBF-dependent gene expression, we examined the dynamics of a fluorescent mVenus-PEST protein expressed by the *CLN2* promoter, a well-established target of SBF ^8,43,44^ (**Fig. 5A**). mVenus was destabilized using the PEST sequence from the *CLN2* gene so that fluorescence more accurately reflects current transcriptional activity^45^. The overall expression patterns of the mutants are comparable as the *CLN2* promoter activates sharply just before bud emergence and peaks about 50 minutes later, consistent with previous results ^6,46^. However, the maximal concentration is reduced by 10-50% (relative to concentration at budding) in several *WHI5* phosphorylation mutants compared to wild-type (**Fig. 5B-D**). The G1 phosphorylation site mutant activates the *CLN2* promoter only slightly less than WT, while the loss of Cdk1 phosphorylation in *WHI5*^Δ*Cdk*1^ cells has a much greater dampening effect on expression, as might be anticipated from its effect on the duration of S/G2/M phases. Interestingly, this effect is not enhanced by increasing protein to wild-type levels in *2xWHI5*^Δ*Cdk1*^ cells, suggesting the S/G2/M expression-inhibiting effect of Whi5 is saturated in these mutants. Transcription can be even further reduced when all phosphorylation sites are removed in the fully unphosphorylatable *WHI5^NP^*mutant. In contrast, *WHI5*^Δ*Cks1*^ cells show negligible difference in expression compared to WT, suggesting that rapid hyperphosphorylation of Whi5 is not necessary for wild type S/G2/M expression dynamics of SBF target genes.

**Figure 5:**
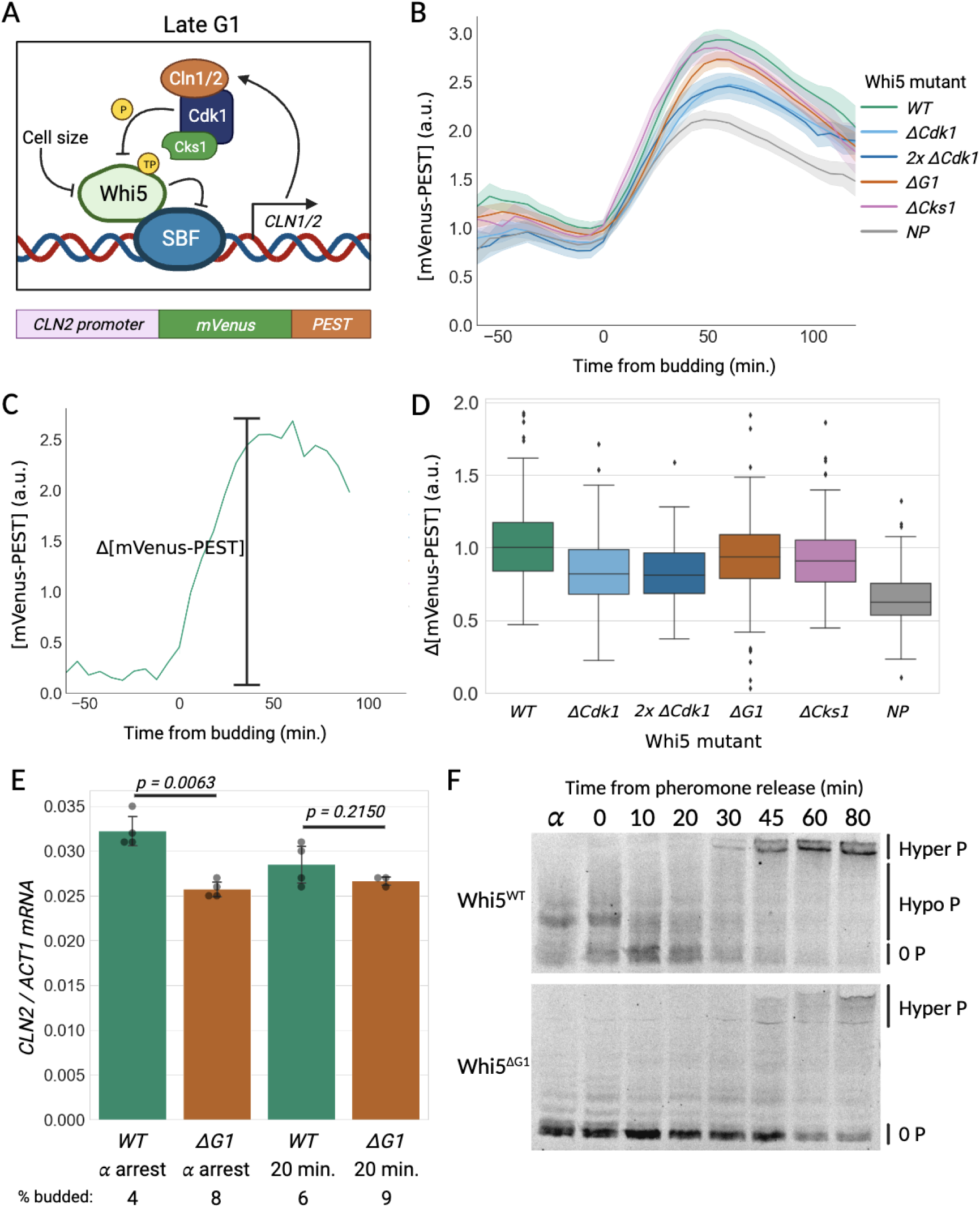
Whi5 phosphorylation regulates expression from a *CLN2* promoter. A) Diagram of Cln1,2-Cdk1 positive feedback regulation with Whi5 and SBF. The *CLN2* promoter driving an mVenus-PEST fluorescent reporter is depicted alongside. B) Concentration of mVenus-PEST driven by a *CLN2* promoter averaged over many daughter cell cycles. Traces are aligned at bud appearance, and scaled such that average mVenus-PEST concentration in WT cells is 1 a.u. at budding. Data lines indicate the mean values and the shaded area denotes 95% c.i. N_WT_ = 230, N_ΔCdk1_ = 207, N_2x ΔCdk1_ = 78, N_ΔG1_ = 251, N_ΔCks1_ = 107, N_NP_ = 101. C) Characteristic single cell trace indicating the amplitude of expression from a *CLN2* promoter over one cell cycle. D) Distributions of amplitudes of *CLN2* promoter activity measured as in panel C for cells of the indicated genotype (same cells as panel B). Box centers indicate median, box edges indicate quartiles, dots indicate outliers (points more than 1.5 times above or below the interquartile range), and whiskers indicate maximum and minimum non-outlier points. Δ[mVenus-PEST] is rescaled such that WT is centered at 1 a.u. E) Mean ratios of *CLN2* to *ACT1* mRNA at the indicated time points, for the indicated mutants, assayed by qPCR. Cells growing in synthetic media with glycerol/ethanol were arrested in G1 using α-factor, then released into the cell cycle through drug washout. Dots represent biological replicates and error bars represent the 95% c.i.; p-values are calculated using a two-tailed Student t-test. The percentage of cells budded from one replicate is indicated at the bottom. F) Phos-tag immunoblot of Whi5-3xFLAG for wild type and ΔG1 variants, per the time course described for panel E. Samples for qPCR were taken from arrested populations and 20 minutes after washout, which is in the middle of G1.

Having assessed peak levels of SBF-dependent gene expression downstream of the G1/S transition, we next sought to examine the level of expression upstream of the transition. The basal level in early G1 is of interest because while small, it likely contributes to initiation of the positive feedback loop that commits cells to division ^35^. However, early G1 levels of *CLN2pr-* driven transcription are low and difficult to detect using fluorescence microscopy. We therefore used quantitative PCR to compare levels of *CLN2* mRNA in cells expressing Whi5^WT^ or Whi5^ΔG1^ early in the cell cycle. Wild-type cells showed significantly higher *CLN2* expression during α-factor arrest than *WHI5*^Δ*G1*^ cells (**Fig. 5E**), correlating with more highly phosphorylated Whi5 (**Fig. 5F**). However, 20 minutes after release into G1 phase, both types of cells showed similar levels of *CLN2* expression, suggesting the hypo-phosphorylation of Whi5^WT^ contributes minimally to Whi5 inhibition in early G1. Our data on post-G1/S SBF target transcription reveal that Cdk1-dependent Whi5 phosphorylation is important for post-G1 *CLN2* transcription to reach a sufficient level to ensure timely progression through S/G2/M. However, the size control effects of mutating G1-specific Whi5 phosphorylation sites are not explained by our pre-G1/S transcription data, suggesting that the phosphorylation of these sites may contribute to size regulation via other targets or other mechanisms.

## DISCUSSION

In this study, we sought to better understand how Whi5 phosphorylation regulates cell cycle progression. Earlier work showed that Whi5 is hyper-phosphorylated at the G1/S transition, most likely by Cdk1-Cln1,2 complexes. However, these studies could not resolve the hypo-phosphorylated state prevalent in early G1 phase when Whi5 inhibits the SBF transcription factor and by extension the G1/S transition. To resolve the Whi5 phospho-isoforms present in early G1, a recent study used Phos-tag gels that specifically retard the migration of phosphorylated residues ^19^. This revealed multiple, likely mono-phosphorylated, isoforms of Whi5 in early G1, which were rapidly hyperphosphorylated at the G1/S transition. This study further showed that this G1 phosphorylation was not due to Cln3-Cdk1 activity, as previously thought. Rather, Cln3-Cdk1’s role in cell cycle regulation is to directly promote transcription at SBF-bound promoters by phosphorylating the C-terminal domain of the RNA polymerase II subunit Rpb1. These results raise several important questions: what are the kinases responsible for Whi5 phosphorylation in early G1, where do they phosphorylate Whi5, and what is the function of these phosphorylations?

To address the function of Whi5 phosphorylation dynamics, we first identified all the phosphorylation sites and their individual phosphorylation dynamics. Our increased resolution using Phos-tag gels allowed us to find a previously unidentified 19th phosphorylation site ^16^, as well as identify the specific sites that were phosphorylated in early G1 (3 Cdk1 sites and 4 non-Cdk1 sites). Mutation of the G1 phosphorylation sites resulted in a delayed G1/S transition, suggesting that Whi5 phosphorylation in G1 promotes cell cycle entry. However, other aspects of the G1/S transition, including Whi5 translocation into the cytoplasm and SBF-dependent transcription, were similar to that in WT cells. Our analysis also revealed that Cks1-dependent priming interactions through phospho-threonines ^26,27^ are essential for the rapid hyper-phosphorylation of Cdk1 target sites on Whi5 at G1/S, although abrogating this docking interaction only had a small effect on cell size regulation.

### Whi5 phosphorylation is required for timely progression through G1 and S/G2/M phases of the cell cycle

Our work highlights the importance of phosphorylation for inactivating Whi5 and driving cell cycle progression. While we found that Whi5 phosphorylation regulates the G1/S transition, as expected, our analysis also revealed an unexpected effect in S/G2/M. Cells expressing Whi5 proteins lacking all Cdk1 sites had an extended S/G2/M phase of the cell cycle, which could be due to Whi5^ΔCdk1^ being retained in the nucleus during S/G2/M and decreasing SBF-dependent expression. It is worth noting that this effect of removing Cdk1 sites decreased Whi5 synthesis so that two copies of the gene were required to recover wild type protein concentrations. This unanticipated effect on synthesis may explain the more modest size phenotypes of Whi5 Cdk1 site mutations reported in a previous study ^16^.

### Whi5 processes multiple signals to regulate G1/S timing

That phosphorylation of Whi5 during G1 affects inhibition of the G1/S transition supports a model wherein Whi5 integrates multiple input signals to control the cell cycle. Consistent with previous results ^21–23,25^, we found that Whi5 concentration reflects the instantaneous size of a cell in all cases we examined. While Whi5 phosphorylation site mutations can affect the synthesis rate of Whi5, they do not impact the general pattern of S/G2/M synthesis followed by dilution in the subsequent G1 phase. In addition to cell size, phosphorylation modulates the overall inhibitory activity of Whi5 in G1. The exact molecular mechanism of how loss of phosphorylation reduces SBF target expression remains unclear, but we speculate that different G1 phosphorylation sites reduce Whi5 binding to SBF and thereby slightly increase SBF-dependent transcription prior to G1/S. Further work is necessary to determine the kinases responsible for phosphorylating the 7 sites targeted in early G1. This is likely to be multiple kinases since the 7 G1 sites are associated with both Cdk1 and non-Cdk1 targeting motifs.

### G1/S phosphorylation dynamics are conserved from yeast to human

Budding yeast has long been used as a model organism for studies of the eukaryotic cell cycle. Both yeast and human cells feature an upstream cyclin complex (Cln3-Cdk1 or Cyclin D-Cdk4/6), followed by a positive feedback loop based on the inhibition of a transcriptional inhibitor (Whi5 or Rb) and a rapid increase in cyclin-dependent kinase activity (Cln1,2-Cdk1 or Cyclin E,A-Cdk2). This in turn relieves inhibition of a transcriptional activator (SBF or E2F) to drive expression of genes necessary for cell cycle progression. However, while there are many parallels between the yeast and human cell cycle networks, the molecular details differ substantially ^47^. For example, E2F and Rb share no evolutionary history or sequence similarity with SBF and Whi5, respectively. Despite this, the molecules follow similar cell-cycle-dependent dynamics as Rb and Whi5 are both diluted in G1 phase to link cell growth to cell division ^22,48^. Their phosphorylation dynamics are also similar: Whi5 and Rb are both hypo-phosphorylated in early G1 and then inactivated through hyper-phosphorylation at the G1/S transition ^16,49^.

The observation that Rb is hypo-phosphorylated (or mono-phosphorylated) in early G1 has led some to question its role as a cell cycle activator ^49,50^. This is because hypo-phosphorylated Rb still acts as an inhibitor of E2F-dependent transcription, which is only fully activated after the G1/S transition following Rb *hyper*-phosphorylation. This led to the hypothesis that Rb hypo-phosphorylation actually serves to activate Rb’s ability to inhibit E2F ^49^ and to form different protein-protein interactions ^51^. However, mutation of Rb’s C-terminal helix that docks cyclin D, which presumably results in loss of Rb mono-phosphorylation, makes Rb a *more* effective inhibitor of the cell cycle ^52^. Thus, mono-phosphorylation of Rb by cyclin D could promote cell cycle progression, although it is unclear how this might work at a molecular level.

The conservation of systems-level features of eukaryotic G1/S regulation suggests that Whi5 phosphorylation dynamics in yeast may inform studies of the parallel dynamics of Rb in human cells. Here, we found that mutations removing the G1 phosphorylation sites of Whi5 result in a stronger cell cycle inhibitor and reduced SBF-dependent gene expression. This supports a model in which the hypo-phosphorylation of Whi5 and Rb in early G1 results in their weak inactivation and slightly increased activity of their targets SBF and E2F. Mild expression of SBF and E2F targets such as downstream cyclins Cln1,2 and Cyclins E and A, in concert with the continual dilution of the inhibitors Whi5 and Rb, eventually results in activation of a positive feedback loop committing cells to division. Thus, these key inhibitors of cell cycle progression may serve to integrate cell size sensing with additional signals during G1, providing an avenue for cells to modulate cell cycle control in response to changing intra- and extra-cellular signals. While more work is needed to test this model, particularly in human cells, the extensive conservation of cell cycle dynamics from yeast to humans argues in its favor.

## Acknowledgements

We thank Kurt Schmoller and members of the Skotheim laboratory for discussions and feedback on the manuscript. This work was supported by a Chan Zuckerberg Biohub Investigator Award (J.M.S.), and the NIH (R35 GM134858).

## Author contributions

J.X. and J.J.T. performed the experiments, J.X., J.J.T., M.K., and J.M.S. designed the experiments, and J.X. and J.M.S. wrote the manuscript.

## STAR METHODS

### RESOURCE AVAILABILITY

#### Lead Contact

Further information and requests for resources and reagents should be directed to and will be fulfilled by the lead contact, Jan Skotheim.

#### Materials Availability

All plasmids and strains generated in this study are available from the lead contact upon request without restriction.

#### Data and Code Availability

- Original western blot images have been deposited at Figshare and are publicly available as of the date of publication. DOIs are listed in the key resources table. Microscopy data reported in this paper will be shared by the lead contact upon request.
- All original code has been deposited at Github and is publicly available as of the date of publication. DOIs are listed in the key resources table.
- Any additional information required to reanalyze the data reported in this paper is available from the lead contact upon request.

### EXPERIMENTAL MODEL DETAILS

Standard procedures were used for growth and genetic manipulation of *Saccharomyces cerevisiae*. Full genotypes of all strains used in this study are listed in Table S1 and the Key Resource Table. All strains used were derived from the W303 background. Cells were grown at 30°C in yeast extract/peptone medium with 2% glucose (YPD), or in synthetic complete medium with 2% glycerol and 1% ethanol (SCGE).

### METHOD DETAILS

#### Cell size measurements

Cell volume was measured using a Beckman Coulter Z2 particle counter (Beckman Coulter). Cells were grown to log-phase (OD600 0.08-0.2) in SCGE media then sonicated briefly. 80-150 µL of sonicated cells were diluted into 10 mL of Isoton II diluent (Beckman Coulter #8546719). Two measurements of 10,000-20,000 each were averaged for a final measurement per sample. Particles below 10 fL and over 300 fL in volume were excluded from analysis.

#### Mating pheromone cell cycle synchronization

For protein extraction and Phos-tag analysis, cells were grown to log-phase in 100-150 mL YPD media (∼90 minute doubling time) then arrested with the addition of α-factor (1 µg/mL, Peptide 2.0, Chantilly VA) for 150 minutes (∼1.6 times generation time). Arrest was confirmed by microscopy analysis checking for shmoos. Cells were spun down in a tabletop centrifuge (939 g, 1 minute) and resuspended in an equal volume of fresh media three times to wash out α-factor. 15 mL of cells were collected before washout (t = α), immediately after washout (t = 0), and at indicated time points thereafter. For collection, cells were spun down (939 g, 2 minutes), media was removed, then pellets were flash-frozen in liquid nitrogen.

For RNA extraction and qPCR, cells were grown to log-phase in 10-30 mL SCGE media (∼150 minute doubling time), then arrested using α-factor (1µg/mL, Peptide 2.0, Chantilly VA) for 240 minutes (∼1.6 times generation time). Arrest was confirmed by checking for shmoos under light microscopy. Cells were washed out using a vacuum-driven filter (0.22 µm PES membrane, GenClone) and resuspended in an equal volume of fresh media three times to wash out α-factor. 1.5 mL of cells were collected before washout (t = α), and at indicated time points thereafter as labeled in Figure 5E. For collection, cells were spun down, media was removed, then pellets were flash-frozen in liquid nitrogen. At each time point, 450 µL of culture was also collected and fixed with 50 µL of 37.5% formaldehyde. Fixed cultures were visually inspected to determine the percentage of cells budded. 100-200 cells were counted for each sample. 450 µL of culture was also added to 1 mL 100% ethanol to fix for DNA staining and flow cytometry.

#### Protein extraction and Phos-tag gel electrophoresis

To extract protein, frozen cell pellets were resuspended in urea lysis buffer (20 mM Tris•Cl pH 7.5, 7 M urea, 2 M thiourea, 65 mM CHAPS, 65 mM DTT, 50 mM NaF, 80 mM beta-glycerophosphate, 1 mM sodium orthovanadate, 1 mM PMSF) then lysed by bead beating using an MP Bio FastPrep-24 (MP Biomedicals, Santa Ana, CA; settings: 5.0 m/s, 1 x 40 seconds) in a cold room at 4°C. Cell debris was pelleted by centrifugation (17,000 g, 10 minutes) and the supernatant recovered.

Protein lysates were separated on tris-glycine or tris-acetate SDS-PAGE gels and transferred to a nitrocellulose membrane using the iBlot 2 dry blotting system (Invitrogen #IB21001). To separate different phospho-Whi5 species, collected protein lysates were separated using 10% SDS-PAGE supplemented with 50 or 75 μM Phos-tag reagent (Wako Pure Chemical Industries). Protein amount was estimated using the Bradford assay (5000006, Bio-Rad, Hercules, CA) and approximately 10 μg of protein was loaded per well, along with 6 μL of Laemmli sample buffer. Total volume was equalized between samples using urea lysis buffer. Western-blotting was done using anti-FLAG (mouse, monoclonal, Sigma-Aldrich Cat#F3165) at 1/1,000 dilution. Primary antibody was detected using IRDye 800CW Goat anti-Mouse (Licor) fluorescently labeled secondary antibody at 1/10,000 dilution. Membranes were then imaged on an Amersham Typhoon 9210 (GE Healthcare Life Sciences) or LI-COR Odyssey CLx scanning machine.

#### Phosphatase assay

50 μL phosphatase reactions were prepared using NEB lambda phosphatase (P0753, NEB, Ipswich, MA). 1.5 μL of high-concentration cell lysate was used in each reaction, and reactions were supplemented, as indicated in the figures, with PhosSTOP phosphatase inhibitor cocktail (04906845001, Roche, Mannheim, Germany). Reaction mixes were incubated at 30°C for 20 minutes, after which 10 μL 6X Laemmli sample buffer was added and the resulting mixture was incubated at 95° for 5 minutes. 15 μL of each sample was loaded into Phos-tag gels for analysis.

#### Protein purification and *in vitro* phosphorylation assays

Cyclin-Cdk1, Cks1, and Whi5 purifications and in vitro kinase assays were performed as described by Kõivomägi et al., 2011 ^19,31^, PMID: 34085218 and Kõivomägi et al., 2021 ^19,31^. In brief, 3FLAG-Cln2-Cdk1 was purified according to published protocols previously used for 3HA-Cln2-Cdk1 complexes (McCusker et al., 2007 ^53^). Cks1 was purified as in PMID: 10913169. 6xHis-tagged Whi5 (wild-type and ΔCks1) proteins were purified by cobalt chromatography and eluted using 200mM imidazole.

For phosphorylation assays, substrate concentrations were kept in the low uM range and the assay conditions were optimized to detect hyperphosphorylated Whi5 species at the end of the time course. Reaction aliquots were taken at indicated time points and the reaction was stopped with SDS-PAGE sample buffer.

The basal composition of the *in vitro* kinase assay mixture contained 50mM HEPES, pH7.4, 150 mM NaCl, 5mM MgCl 2, 20mM imidazole. 0.01mg/ml FLAG peptide, 2% glycerol, 0.05 mM EGTA, 0.2 mg/ml BSA, 500 nM recombinant Cks1 and 500uM ATP (with 2 µCi of [γ-32P] ATP added per reaction (PerkinElmer cat# BLU502Z250UC)]. Phosphorylated proteins were separated on 10% SDS-PAGE supplemented with 50 μM Phos-tag reagent (WakoPure Chemical Corporation) and visualized using autoradiography (Typhoon 9210; GE Healthcare Life Sciences).

#### Live cell microscopy

Cells were grown to early log-phase (OD600 0.08-0.2) in SCGE media, then lightly sonicated before being loaded into a CellASIC Y04C microfluidics plate (Millipore SIGMA) under continuous SCGE media flow at 2 psi. Unless otherwise specified, images were taken every 6 min using an Observer Z1 microscope equipped with an automated stage, a plan apochromat 63x/1.4NA oil immersion objective, and an Axiocam 705 mono camera.

Whi5-mCitrine and mVenus expressed from a *CLN2* promoter were imaged using 400 ms exposures under illumination from a Colibri 505 nm LED module at 25% power. Myo1-3xmKate2, used to determine timing of budding, was imaged by exposure for 1000 ms using a Colibri 555 nm LED module at 25% power.

Cells grown in the microfluidics plate were checked for their average growth rate and experiments where the average cell growth rate deviated from the expected growth rate for those conditions in liquid culture were excluded. For all strains, at least two biological replicate experiments (i.e. separately inoculated cultures) were performed.

#### Flow cytometry

Cells were grown to early log-phase (OD600 0.08-0.2) in SCGE media, then lightly sonicated. 1 mL of cells was added to 1 mL 100% ethanol. Fixed cells were then pelleted and resuspended in sodium citrate (50mM, pH 7.5). Cells were then treated with 0.25mg/ml RNase A (1 hour, 55°C) followed by 10 µL of 20 mg/mL Proteinase K (1 hour, 55°C). Finally, DNA was stained with 10µl of 1.6mg/ml propidium iodide (overnight in the dark, 4°C). Samples were vortexed then measured on an Attune NxT cytometer. Data was analyzed in FlowJo. For all strains, at least two replicate experiments (*i.e.*, separately inoculated cultures) were performed.

#### RNA extraction and qPCR

To extract RNA, frozen cell pellets were resuspended in urea buffer then lysed by bead beating using an MP Bio FastPrep-24 (MP Biomedicals, Santa Ana, CA; settings: 5.0 m/s, 1 x 35 seconds) in a cold room at 4°C. Cell debris was pelleted by centrifugation (17,000 g, 10 minutes) and the supernatant recovered. RNA was then extracted using a Direct-Zol RNA microprep kit (Zymo Research). Complementary DNA was synthesized using the SuperScript IV VILO synthesis kit (Thermo Fisher Scientific). qPCR reactions were performed using iTaq Universal SYBR Green Supermix (Bio-Rad). For each quantified time point, at least three biological replicate experiments (*i.e.*, separately inoculated cultures) were performed.

### QUANTIFICATION AND STATISTICAL ANALYSIS

#### Live cell microscopy data processing

Data from live cell microscopy were first corrected for non-cell background intensity before being processed for segmentation and tracking in the Cell-ACDC analysis framework (Padovani et al., 2022 ^54^).

For quantified fluorescence channels, raw images were first subtracted for dark-field intensity (*i.e.*, signal present even in absence of illumination), then rescaled pixel-by-pixel to a mean image with no cells to account for reproducible spatial variations in the imaging area. Cells were then segmented by phase-contrast images using the YeaZ neural network (Dietler et al., 2020 ^55^), and total fluorescence calculated as the sum of pixel intensities within the segmented area minus the median non-cell pixel intensity in the same image. Fluorescence totals were then corrected for autofluorescence by subtracting a volume-dependent autofluorescence amount obtained from a linear fit to data from cells expressing no signal in the relevant channel.

Geometric volume was estimated by performing a rotation of the segmented area around the longest axis. Mother-bud pairs were assigned in Cell-ACDC, using appearance and disappearance of Myo1-3xmKate2 to determine timing of budding and division respectively. For localization of Whi5-mCitrine, the nuclear localization metric is the coefficient of variation of Whi5-mCitrine signal over the cell area rescaled to the value in early G1. The coefficient of variation over cell area is an effective proxy for a nuclear tag. Unless otherwise specified, only cell traces from first-generation daughter cells with complete cell cycles were included in presented data.

#### General statistical analysis

The definition of error bars is provided in figure legends. For microscopy data, sample size *N* indicates the number of cells measured, and 95% confidence intervals are bootstrapped. All regression is performed on original data; binning of scatterplots (such as in Figure 4) is only for plotting purposes.

---

**Supp. Figure 1.**
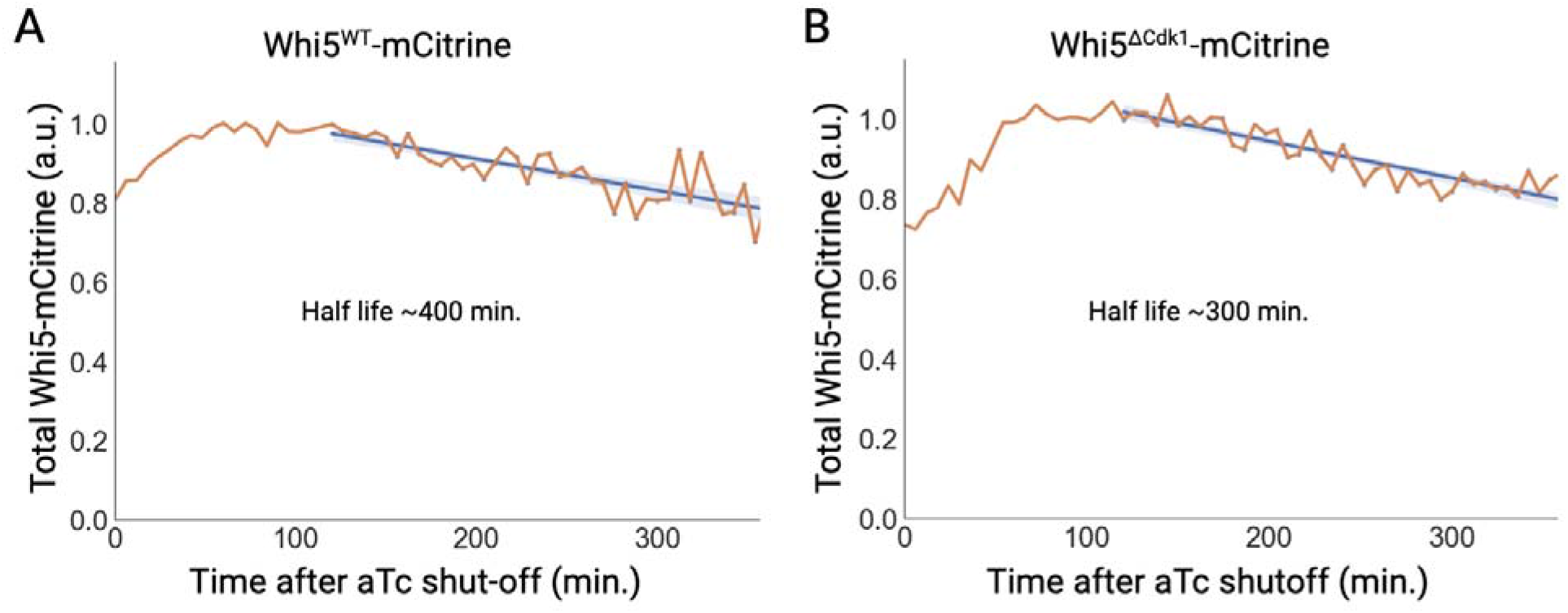
A) Total Whi5 fluorescence over time in strains expressing Whi5^WT^-mCitrine from the WTC_846_ promoter, which is induced by anhydrotetracycline (aTc) ^39^. Cells were grown in synthetic complete media (SCGE; generation time ∼150 minutes) with 2 ng/uL aTc for six hours, then given fresh media with no aTc for the remaining six hours of imaging. The orange line indicates the total fluorescence signal across all cells imaged (*i.e.*, cumulative Whi5-mCitrine expression in all cells). The blue line represents a linear regression to the total fluorescence beginning at two hours after aTc shut-off to ensure time for residual aTc to wash out. The shaded area represents the 95% c.i. B) The same as A, but with cells expressing Whi5^ΔCdk1^-mCitrine.

**Supp. Figure 2.**
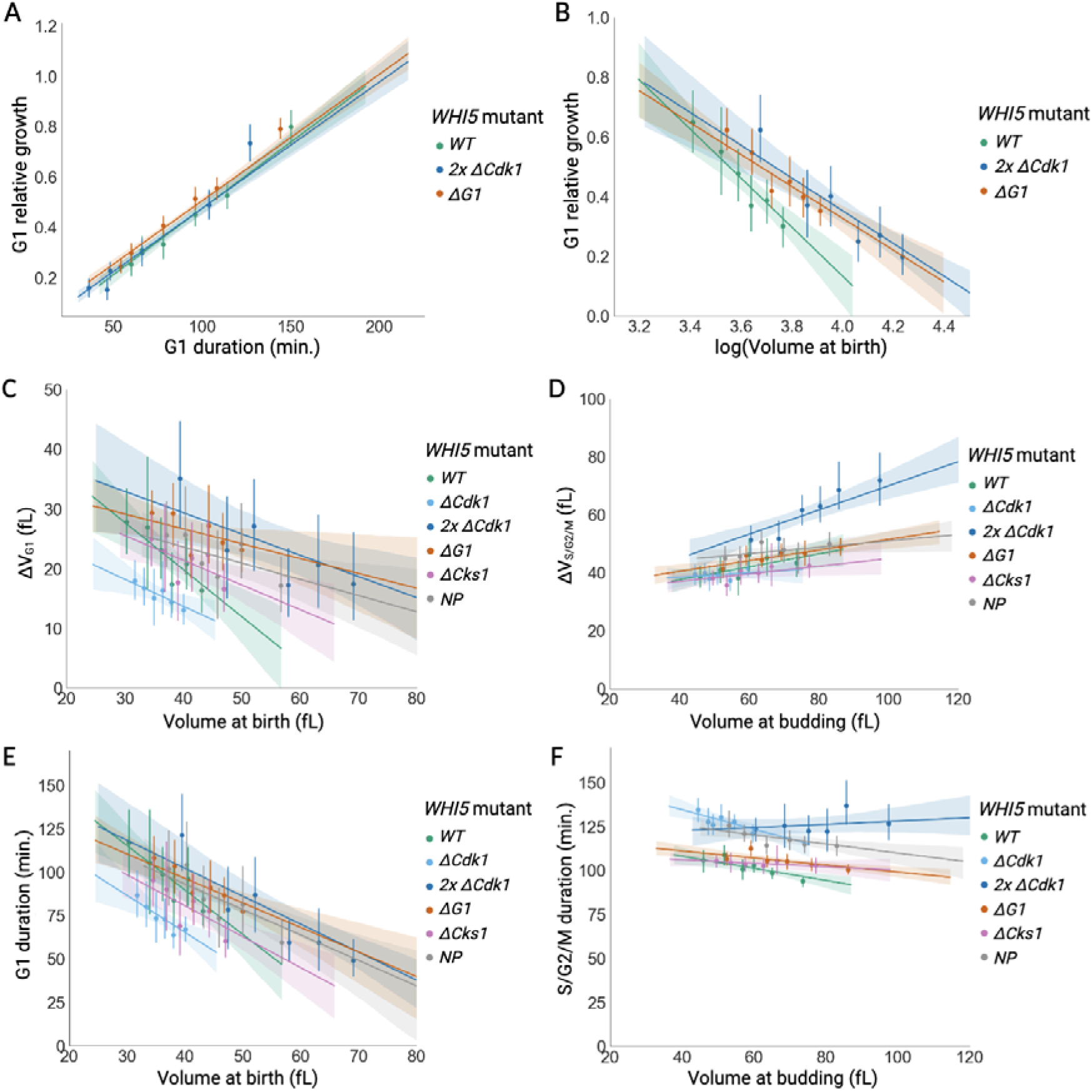
A) Duration of G1 phase versus volume at birth for the same cells as Fig. 4E-H (N_WT_ = 100, N_2x ΔCdk1_ = 98, N_ΔG1_ = 217). Cells are separated into bins with equal numbers. Dots indicate the mean values per bin and error bars denote the 95% c.i. Solid line indicates linear regression to unbinned data and shaded areas denote 95% c.i. B) Ratio of volume at budding to volume at birth, versus volume at birth, for the same cells as Fig. 4E-G. C-F) The same plots as Figure 4E-H, showing all genotypes. N_WT_ = 100, N_ΔCdk1_ = 200, N_2x ΔCdk1_ = 98, N_ΔG1_ = 217, N_ΔCks1_ = 104, N_NP_ = 83.

**Supp. Figure 3.**
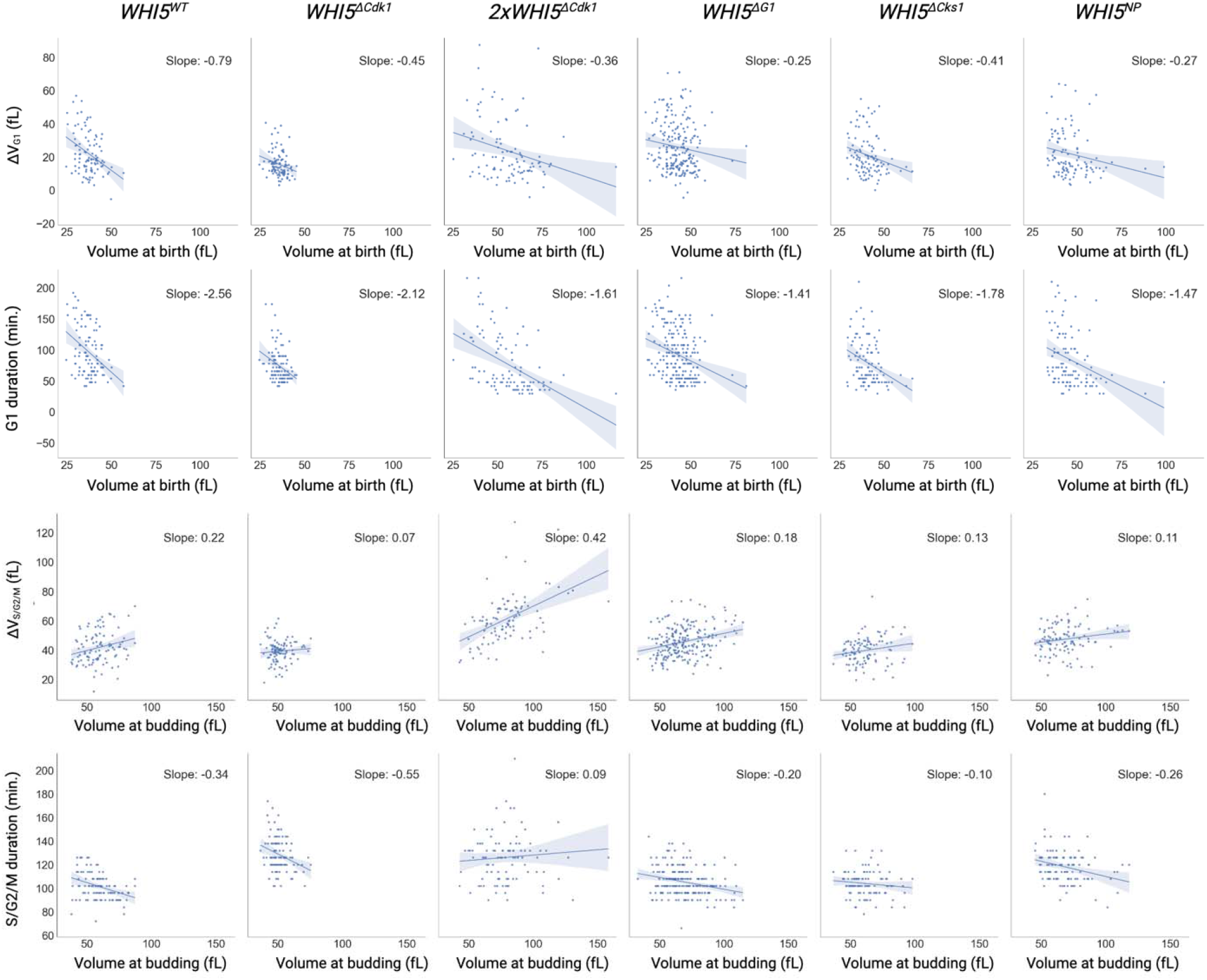
The same data as Fig. 4E-H and S1C-F, separated by *WHI5* phosphorylation mutant. Each point represents a single first-generation daughter cell cycle. Solid lines indicate linear regressions to data and shaded areas denote 95% c.i. N_WT_ = 100, N_ΔCdk1_ = 200, N_2x ΔCdk1_ = 98, N_ΔG1_ = 217, N_ΔCks1_ = 104, N_NP_ = 83.

**Supp. Table 1.**
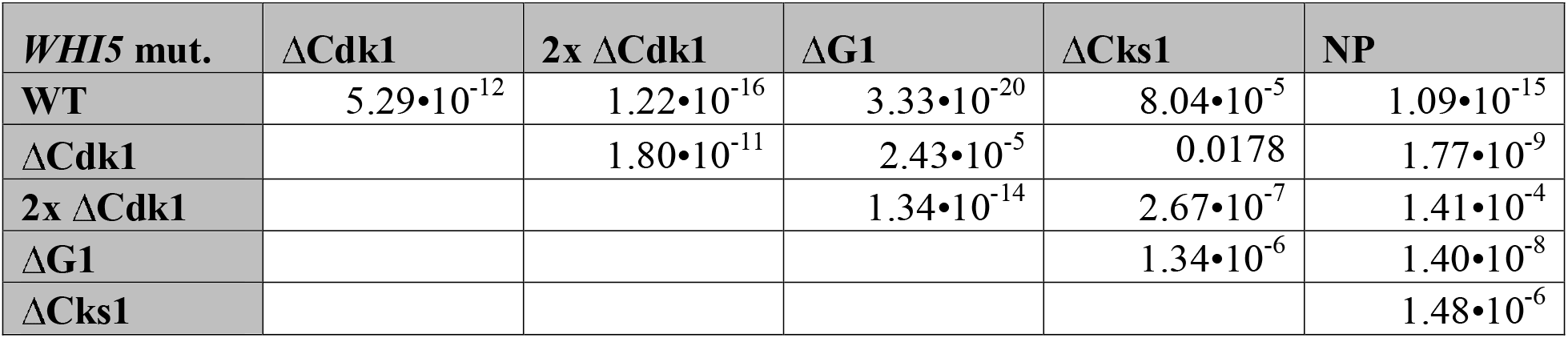
P-values for differences in mean cell size based on Coulter counter data shown in Figure 4C. P-values are calculated using a two-tailed Student t-test.

**Supp. Table 2.** P-values for differences in Δ[mVenus-PEST] shown in Figure 5D. P-values are calculated using a two-tailed Student t-test.

| **WHI5 mut.** | **ΔCdk1** | **2x ΔCdk1** | **ΔG1** | **ΔCks1** | **NP** |
| --- | --- | --- | --- | --- | --- |
| **WT** | $2.53 \cdot 10^{-13}$ | $1.56 \cdot 10^{-8}$ | $1.68 \cdot 10^{-4}$ | 0.00885 | $2.44 \cdot 10^{-29}$ |
| **ΔCdk1** | | 0.815 | $3.48 \cdot 10^{-5}$ | $4.50 \cdot 10^{-4}$ | $3.21 \cdot 10^{-11}$ |
| **2x ΔCdk1** | | | $8.76 \cdot 10^{-4}$ | 0.00195 | $1.03 \cdot 10^{-8}$ |
| **ΔG1** | | | | 0.795 | $1.13 \cdot 10^{-21}$ |
| **ΔCks1** | | | | | $6.24 \cdot 10^{-17}$ |

## Notes

### Competing Interest Statement

The authors have declared no competing interest.

